# Individual difference in social learning in a granivorous passerine bird, and its independence from personality

**DOI:** 10.1101/2025.03.05.641444

**Authors:** Andrea Frascotti, Ricardo Silva, Paulo Gama Mota

**Affiliations:** Universita degli Studi di Milano; CIBIO; Universidade de Coimbra; CIBIO

**Keywords:** Boldness, Memory, Personality, *Serinus serinus*, Sociability, Social learning

## Abstract

Animals use learning to adaptively adjust their behaviour to conditions taking advantage of previous experiences. While individual learning is advantageous, it includes risks and costs that can be overcome by learning from others. Despite its relevance, the ability to learn from observing others’ behaviour was tested in only a limited number of bird’s species, of which only a few feeds mainly on plants. There is still much to be learn what kind of information birds are interested and capable of gathering from conspecifics in the foraging context. We set out to test the social learning skills of a granivorous gregarious species of cardueline finch, the Serin (*Serinus serinus*), because they likely follow conspecifics cues to forage. We used an observer-demonstrator paradigm where observers were given the opportunity to learn to reach hidden food from observation of demonstrators that were previously trained to perform the task. Almost half of the birds tested were capable of learning (40%) from a conspecific in a colour-food association experiment, and learners were able to remember the association after fifteen days. Also, almost half of the birds tested for this revealed to be capable of reversal learning. The ability to learn was not influenced by sex or age, of both demonstrators and learners, but learners had longer wings than non-learners. We investigated whether personality could explain the differences in learning ability of these birds. We assessed individual boldness and sociability through the novel object test and mirror personality tests respectively. Although we found repeatability those personality traits we found no association with social learning ability.

## Introduction

Learning through personal experience is a fundamental process of interacting with the environment and is widespread in animals. However, it exposes individuals to many threats and costs (Kendall et al., 2009), that can be overcome if individuals acquire information from observation or interaction with other animals (typically conspecifics) or its products (Heyes, 1994). In the last few decades many studies on social learning have broadened our understanding of this behaviour in an increasing number of animals with different social lifestyles and neurobiological complexity, from mammals (Thornton & Malapert, 2009) and birds (Templeton and Giraldeau, 1996) to insects (Mery *et al*., 2009) and molluscs (Fiorito and Scotto, 1992).

Birds benefit from social learning in many different contexts such as in migratory routes, predator and prey recognition, mate choice and song learning (Slagsvold and Wiebe, 2011; Hultsch & Todt, 2004). Social learning is also important in foraging. It has been investigated in a variety of cases, such as, how novel information spreads across social networks (Aplin et al, 2013; 2015), the use of social cues to locate food (Katsnelson et al, 2011), the influence of conspecific preferences when faced with a novel food source (Cadieu; Fryday et al, 1994; Benskin et al, 2002), or the learning of hunting strategies from older conspecifics (Kitowski, 2009). Still, there is much to be understood on how the social learning process works and what is exactly the information birds are gathering from conspecifics. Our study is tailored to understand if a small passerine is capable of socially learn a stimulus-response association and investigate why interindividual differences exist in this behaviour.

Social learning may appear to be always an adaptive shortcut to acquire valuable knowledge, but, actually, mindless copying is not an effective strategy. As predicted from theoretical models and observed in nature, animals do not rely solely on indiscriminate social learning (Kameda & Nakanishi, 2002). Instead, natural selection has favoured the establishment of several “social learning strategies” (Laland 2004; Kendal *et al.,* 2005; Kendal *et al.,* 2009). For instances, social learning is preferred when individual learning is costly or when personal information is not updated to the current situation, exemplified in nine-spined sticklebacks and worker honeybees (Van Bergen *et al.,* 2004; H Grüter’s et al., 2010; Grüter & Ratnieks, 2011). It is also important to learn from a successful individual instead of a subordinate one (Greene, 1987), or to copy the majority (Pike & Laland, 2010).

Learning relies on memory in order to benefit from the useful information gathered. In an ever-changing environment, it is essential to adapt previous knowledge to novel circumstances. In some cases, it requires to abandon an association established and replace it by a new one. This cognitive ability can be discerned in tests of reversal learning. Variations of this test have been explored to great success on a variety of mammals, such as rats, macaques, and pigs, (Guillamón et al. 1986; Ha et al. 2011; Roelofs et al. 2017), and also birds, such as corvids and great tits (Cauchoix et al., 2017; Bond et al., 2007). In many cases, individuals were able to learn an associative rule and then, after being exposed to a situation where the rule was reversed, they were capable of adapting their behaviour to the new conditions. Furthermore, some evidence suggests that in guppies and other vertebrates, females are faster and more precise at reversal learning than males (Petrazzini et al., 2017; Guillamón et al. 1986; Ha et al. 2011; Roelofs et al. 2017).

In many species, individuals have been observed to consistently show a preference between personal or social information (e.g. Dubois et al., 2012; Hämäläinen, 2017; Rendell et al., 2011; Rosa et al., 2012). This individual variation in the use of social learning may be related to some individual’s characteristics, such as sex (Rendell et al., 2011; Rosa et al., 2012), age (Thornton & Malapert, 2009) and even personality (e.g. Kurvers et al., 2010; Marchetti & Drent, 2000; Nomakuchi, Park, & Bell, 2009) but tends to vary between species.

Biologists now define personality as an individual behavioural difference that is consistent over time and across different contexts (Rèale *et al*., 2010). Thus, such individual differences may impose constraints and affect the cost/benefit trade-offs associated with personal and social information use. Although a recent review found evidence for a small but significant relationship between cognition and personality (Dougherty and Guillette, 2018), the pattern of association was highly variable, meaning that across the analysed species, when it occurred, personality appeared associated with cognition in different ways. This finding does not align with the prior favoured hypothesis that considers “proactive”, bold/explorative, animals capable of experiencing more of their environment more quickly, thus encountering to-be- learnt associations more readily than “reactive”, shy/less explorative, individuals (Sih and Del Giudice, 2012).

For the less studied case of social learning, the association with personality is also not obvious. Studies performed on great tits and sticklebacks have shown that fast exploring individuals make greater and faster use of social information gained from observing demonstrators (Marchetti & Drent, 2000; Nomakuchi *et al.,* 2009). Similarly, in chacma baboons, bolder individuals were more proficient social learners (Carter *et al*. 2014). By contrast, a subsequent study also with great tits found the opposite, in that slow explorers based their behaviour on prior knowledge of both personal and social origin more than fast explorers, who tended to act less according to available information (Smit & van Oers, 2019). Other similar findings, where bolder individuals prefer to ignore or even avoid conspecifics, were reported for guppies (Trompf & Brown, 2014) and barnacle geese (Kurvers *et al*., 2010).

Social learning has been documented in a number of bird species, especially active hunters such as great tits, while less is known about social learning on birds that feed most or exclusively on plants. Furthermore, it is still unclear what kind of information birds are interested and capable of gathering from conspecifics in the foraging context. We, therefore, designed an experiment to know if a complex associative learning task could be learnt socially in a foraging context by serins (*Serinus serinus*). Serins are granivorous finches that feed almost entirely on a short selection of seeds and plant parts (Diaz, 1994) and are unique in that they exclusively feed their young with seeds (Valera et al., 2005). Serins gather in flocks, which are particularly large outside the breeding season, to forage communally on seeds and other plant parts that are distributed in patches across the farmland. Here social learning could be beneficial not only to locate food but also to learn how to exploit novel food sources. Serins, along with the other cardueline finches, have not yet been studied on their cognitive skills.

Following previous findings on gregarious species of birds, we expected that social learning would vary among individuals, with just part of them revealing the ability to learn fast (Aplin et al., 2013; 2015; Kurvers et al., 2010). In addition, we hypothesised that traits such as the age and sex of the observers might have a role in the social learning capabilities of the birds. We set a colour-food association task that involves observers and demonstrators to test if serins are capable of learning by observing the behaviour of others, and if social information use is dependent on sex, age, or physical characteristics of the birds. We also wanted to further investigate the learning skills of these birds by testing their retention memory and the ability for reversal learning. Finally, considering the existence of individual differences in social learning, we investigated whether they resulted from differences in personality. We expected more sociable observers to be more attentive of the demonstrator performance and thus increasing their chance of social learning. Bolder individuals were expected to interact more easily with the foraging board giving them the possibility to perform the observed behaviour from the beginning, without being driven away from novelty.

## Materials & Methods

### Subjects and housing

Wild serins were captured in 2021 and 2023, in rural areas around Coimbra (Portugal). A total of 22 serins, 11 female and 11 males, were captured in 2021 and 25, 14 females and 11 males in 2023. All 47 adults captured were transported to the aviary of the University of Coimbra, where they were housed by sex in cages,100 x 50 x 50 cm in size, holding a maximum of four individuals and kept under natural light. Cages were provided with water and food ad libitum containing a standard mix of seeds for passerine birds, with supplementation of vitamin b and calcium that was provided as necessary. The testing room was mostly illuminated with artificial light emitting a light spectrum similar to sunlight.

All testing sessions were filmed with two cameras, one framing the whole testing area and one zooming into the testing apparatus. Filming and behavioural scoring were made using the software Observer XT 10. All birds were ringed with plastic black numbered rings (AvianID), and several measurements were collected from each individual: age, weight, left and right wing length, tarsus length, beak length, beak width, beak height and ecto-parasite load (External Dataset). Only two age classes could be considered: first year and older birds, which was based on the moulting of the coverts wing feathers (Svensson, 1992). Morphometric variables were normally distributed (Shapiro-Wilkins test) and were included in a Principal Components Analysis (PCA) to reduce the seven morphometric variables into two Principal Components that explained 60% of the variance. PC1 had positive loading on wing length (Table A.1), while PC2 had positive loading on the height and width of the beak and negative on beak length. The birds were released in their capture sites by the end of the study.

### Colour-food association task

The social learning test consisted of a foraging task in which birds were rewarded with hidden food when they displaced the disks of specific colours. Birds were divided in two role groups demonstrators and observers. The first ones were trained to learn a food-colour association through a gradual learning process. The latter had the opportunity to learn the association from the observation of the demonstrators’ behaviour. The food-colour association task was constituted of a wooden board (Fig. 1) with sixteen wells arranged in a 4x4 matrix. During tests half of the wells were covered with plastic disks of two colours: four red and four blue. Disk colours were chosen for high contrast and for not being part of the birds’ colouration. Food was only available in the wells covered with the colour the demonstrators were trained with. Both during training of demonstrators and the trials of the observers, the colour disks were distributed over the wells in a randomized way in each trial by applying random permutations generated in R v.4.3.1 (R Core Team, 2023). A small amount of seeds was placed in the wells covered by the disks of the association colour. In order to have access to the food, the birds had to displace the disks covering it. The colour-food association remained consistent across all trials for the same bird. The randomization of disks location prevented any possible association between food and position. After each trial, the disks and the board were washed to exclude any possible olfactory cues.

**Figure 1:**
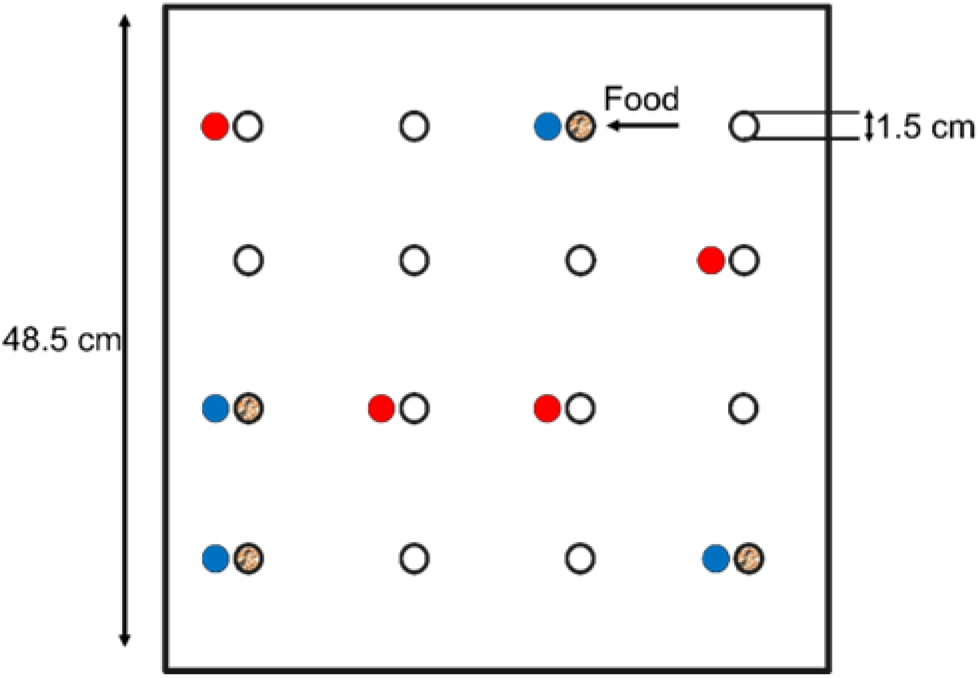
Graphical representation of the foraging task. The square represents a wooden board with sixteen equally distant wells carved on the top side. Eight disks of two colours are placed in correspondence of the wells with a random disposition in each trial. For each bird food is present in the wells associated with only one of the two disks’ colour.

### Demonstrators training

Ten birds (5 males and 5 females; 5 first year and 5 older) were randomly chosen to be trained as demonstrators. Of these, only seven (4 females and 3 males; 3 first year and 4 older) were caable of learning following the defined criterion. Demonstrators underwent 3 hours of fasting before every training session. In each trial the demonstrator was moved into the testing room and exposed to the novel foraging task for 30 minutes. A 10 minutes break was established between the first and second trial of the day. The training program was developed considering three consecutive phases (Fig. 2): the coloured disks were (*A*) positioned on the side of the wells containing food that was visible to the bird when approaching the well; (*B*) partially covering the wells so that the food inside was only visible at a close inspection; (*C*) fully covering the wells, preventing visual access to the food, so that birds could only access it by displacing the disks. During phases *A* and *B*, the learning criterion for demonstrators under training was that they visited at least three wells containing food in three out of four consecutive trials. Only then they proceeded to the next phase. In the final phase (*C*), the learning criterion was to visit at least three wells of the right colour that contained food, but not a single well of the wrong colour, in three out of four consecutive trials. If one individual did not reach the learning criterion after 25 trials of the same training phase, it was discarded as a possible demonstrator, and a new subject was chosen to start the training. Half of the demonstrators were trained to associate the colour red with the presence of food and the other half the colour blue.

**Figure 2:**
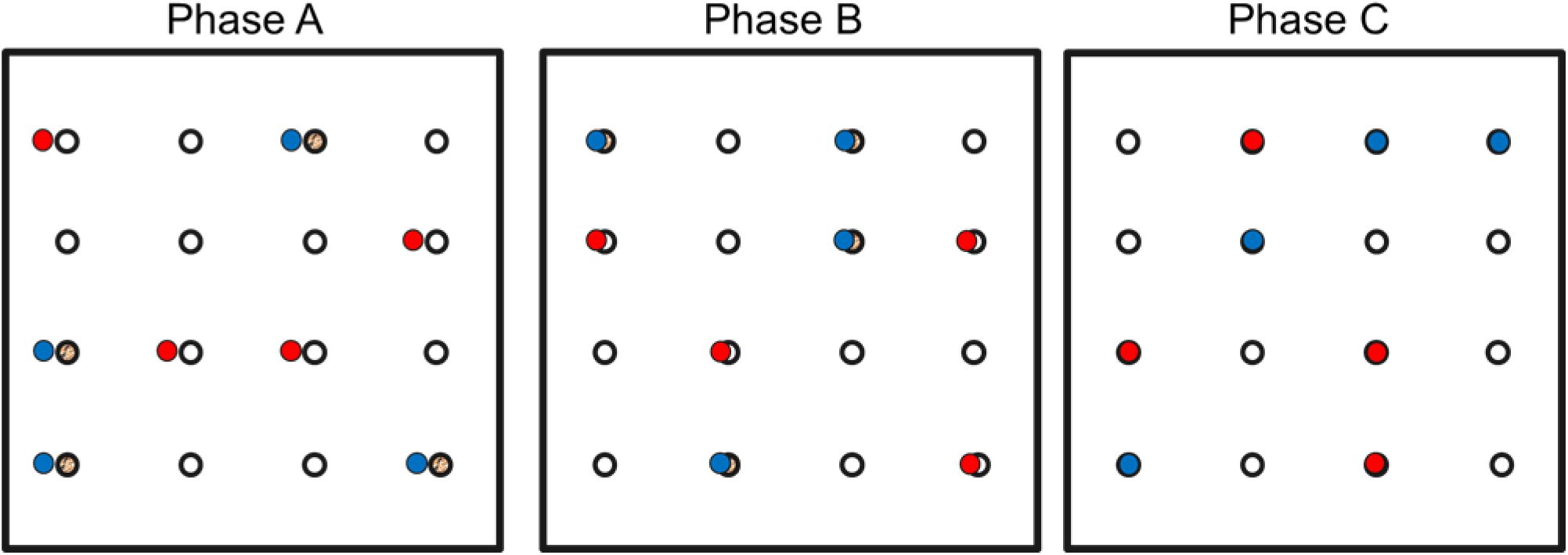
Diagram of the three successive training phases; Phase A) Disks on the wells’ side; Phase B) Disks half cover the wells; Phase C) Disks completely cover the wells.

### Social learning test

In this test, the 35 observers (19 females and 16males; 14 first year and 21 older) were paired with an individual from the demonstrator’s group; pairings were decided to have an even number of the four sex pairs combinations (all pairings are indicated in the external dataset).

At first observers were individually allowed to observe a demonstrator interacting with the foraging apparatus (*observation phase*), and were subsequently released to search for food (*performing phase*) (videos available as supplementary materials). Both the observer and the demonstrator were food deprived for 3 hours prior to the test. This measure ensures a standardized minimum level of motivation and, most importantly, it increases the detection of the learning phenomenon in a foraging context because. In fact, 3h is a reasonable time for birds of this size to make them interested in searching for food. For the *observation phase,* the observer and the demonstrator were brought into the test room. During this *phase* the demonstrator had access to the foraging board while the observer was held in a small cage from where it could observe the demonstrator activity. The observer was kept in the cage to prevent it from distracting the demonstrator from the task. This phase lasted 30 minutes, after which both birds were removed from the testing room, and the board replaced by one with a new disposition of disks. The *observation phase* was only considered a successful learning opportunity - a necessary condition to proceed for the next phase - if the demonstrator visited at least three holes of the right colour and none of the wrong one. If this condition was not met the trial was ended and repeated the next day (this just occurred 3 times over 350+ trials). After a 10 minutes break, the *performing phase* was initiated, with the observer being released in the room and free to move around and interact with the foraging board for 1 hour with no interference. Observer were tested for a maximum of ten trials or until reaching the social learning criterion, that is to have found the food under at least the first three holes visited in at least three trials out of four consecutive ones. The only exception was if a bird after its tenth trial was one away from reaching learning criteria, in which case one extra test was performed. Trials of the same individual were conducted one on every other day to avoid stressing the birds with consecutive food deprivation.

A *control group* of 8 (5 females; 3 males; 4 first year; 4 older) observers was tested in the same way, except that they were exposed to naïve demonstrators, that is, birds which were not trained as demonstrators, in order to determine how likely is that naive birds could learn to meet the learning criterion just by themselves. Achieving it would provide a non-social alternative explanation for learning.

### Memory retention

A subgroup of observers (Subgroup1; n=9) that learnt to perform the social learning test was re-tested 15 days after finishing the social learning task to assess their memory retention ability. These birds were placed in the testing room with the foraging board with a random distribution of disks as in a normal trial, after a 3 hours of food deprivation. Here each bird had the opportunity to eat from the covered holes, and the learning criterion was the same used in the social learning test. The memory test duration was half of a normal trial because all individuals were observed to complete the foraging task in less than 30 minutes, hence the test were not prolonged uselessly to avoid stressing the birds.

### Reversal learning

The Subgroup1 that took part in the memory retention test was subsequently enrolled in a reversal learning test. Here birds were exposed to the foraging apparatus as in the *performing phase* for other 10 trials of 30 minutes each. Only one major difference got introduced in the protocol: the colour-food association previously learnt by each subject was reversed. The criterion to succeed in the trial was the same as in the social learning test. Birds were considered proficient in reversal learning if they succeeded in two trials in a row.

### Personality tests

Since some birds did learn the colour-food association and others did not, we decided to test if this difference between birds could be explained by differences in personality. Thus, a sub- group of birds (Subgroup 2; n=17) were subjected to personality tests in order to determine the presence of some personality traits related with reaction to novelty and sociability. Two standard personality protocols usually used with birds were applied to serins.

### Novel object test

To measure boldness/shyness of birds, we performed a novel object test following a protocol similar to that of Kimball *et al*. (2022). It consists in exposing the subject to the presence of an unfamiliar object placed near the only feeder left in the cage for 15 minutes, after a period of food deprivation of 2 hours by removal of all feeders to standardize motivation (video available as supplementary materials). The protocol comprised five trials per bird, three exposing the subject to different novel objects and two controls without any object. The objects presented were chosen to differ in shapes and colours to avoid habituation (see Supplementary materials), while the order of presentation was randomized. All trials were video recorder to avoid the presence of an experimenter in the room. Trials were carried out between 1100 hours and 1300 hours on different days, over the course of a couple of weeks to assess the stability of the behaviour. The tests were performed in the home cage of each individual, so that the only novelty presented was the object (Fig. 3A). The video recordings were analysed using The Observer XT 10 program. Four variables were considered: the latency to perch near the object, the latency to perch on the feeder and the number of feeding events (each event starts when the bird feed and ends when the bird first leaves the perch), to estimate neophobia, and the number of flights and hops (jump from the ground that travels at least the same distance present between two perches) as a measure of general activity.

**Figure 3:**
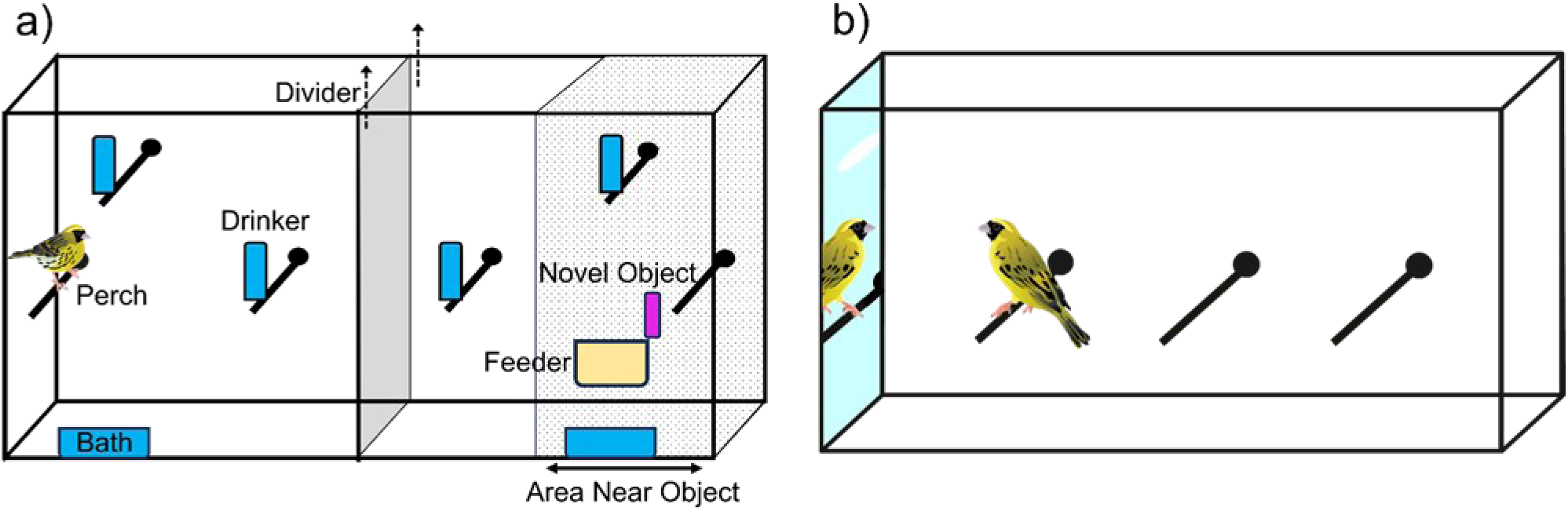
**a)** Graphic representation of the Novel Object Test setup; the Divider was removed at the beginning of the test after a 2h starvation period. **b)** Graphic representation of the Mirror Test setup; the focal bird was placed alone in a cage with a mirror on one side.

### Mirror test

To measure individuals’ sociability, we conducted a mirror test that simulates the presence of an unfamiliar conspecific through the use of a mirror. The test consists in keeping the focal individual alone in a cage with a mirror covering one of the sides of the cage that was initially hidden (video available as supplementary materials). The cage had 3 equally distant perches (Fig. 3B). A two-minute habituation time was established at the start, when the bird was positioned in the cage. The test lasted for 10 minutes, divided in two 5 minutes parts. The mirror was covered by a cardboard during the first half of the test and was exposed by pulling a string to remove the cardboard. Trials were video recorded and analysed as in the novel object test. Six measures were collected in each part of the test: number of fast moves (every movement that travels at least from a perch to another), time spent resting, time spent moving, time spent facing the mirror, mean distance from the mirror and number of pecks on the mirror.

### Statistical analysis

Learning success was tested with a Fischer-Irwin Exact Test to determine if learning could be individual and not social, by comparing the outcomes of experimental and control groups.

Due to the limited sample size to test the effect of different predictors on the social learning results, a series of Generalised Linear Models (GLM) each with one predictor was used instead of a single GLM comprising all predictors. The dependent variable was the outcome of learning with a binary logistic distribution and identity as the link function. The social learning predictors considered in the models were: observer age and sex, demonstrator age and sex, morphology of the observer, a two-way interaction between observer and demonstrator sexes and a two-way interaction between observer and demonstrator and ages. Then a model selection framework was used to compare AICc of the models to better assess evidence for association of these predictors with social learning. Memory ability was tested with a binomial test, the probability of succeeding by chance was 0.5 and the two possible outcomes were “remember” or “do not remember”. The effect of sex and age on reversal learning was tested with two separate GLM due to the small sample.

A recent study demonstrated that blue and great tits, juveniles especially, show an innate bias for the red colour in the foraging context (Teichmann et al., 2020), thus we conducted a simple chi-square statistic between learning result and colour to investigate if serins show a similar bias.

In order to measure the repeatability of the personality variables their distribution had to fall into a known family. The few variables that did not have a known distribution or could not be transformed to reach one were excluded from the analyses. The two latencies measured in the novel object test were very much close to binomial, and were transformed into such, thus describing if the subject did or did not get close to the feeder or the object. Time facing the mirror was square root transformed. General activity and 1/’time resting’ were log transformed. The variables used as predictors in GLM were also tested for correlation. Repeatabilities of the personality variables were tested using the rptR package (Stoffel et al. 2017) for R v4.3.1(R Core Team, 2023).

The control trials of the personality tests were used to determine if the variables did measure a behavioural response to stimuli. For that purpose, a Generalized Linear Mixed Model (GLMM) was performed, with Bird ID as a random factor, and the test nature (control vs experimental) as a fixed factor; a significant effect of the fixed factor would indicate the variable was capturing what was being measured in the personality test.

The personality variables that showed repeatability were aggregated in two PCA, one for each of the personality tests, to obtain an index for subsequent analysis. We used the eigenvalues superior to 1 criterion to retain factors. One principal component was retained from each PCA, which became the Sociability Index and the Boldness Index. These indexes described each bird location on the personality spectrum and were used to test whether personality could explain the outcome of social learning, through a GLM with learning as a binary logistic dependent variable, as above, and the personality traits as predictors.

### Ethical note

All experimental procedures conducted on the birds comply with Portuguese and European laws regarding scientific testing on animals. The capture of birds and their subsequent maintenance for the experiments of this project were carried out under permits N°09/2021/CAPT and N° 049/2023/CAPT to (Author name), by ICNF, Instituto da Conservação da Natureza e das Florestas, I.P. Research was also assessed and approved by the body responsible for the well-being of animals at CIBIO-BIOPOLIS (2024/03). No bird was subjected to any act of mistreatment or neglect. Sampling was conducted during the nonbreeding season making use of fine-mesh mist nets that were inspected every 20 minutes to minimize the harmfulness of the process. All birds were looked after and monitored daily by the authors of this work throughout their seven-month captivity period. To minimize distress a limit of 4 hours of food deprivation per day was set, also individuals that partake in aggressive interactions were moved to different cages to prevent future ones. At the end of the experimental procedures, all birds were released in the same locations where they were captured.

## RESULTS

### Social learning

Of the 35 birds that were tested after observing experienced demonstrators visiting the holes covered with the appropriate colour disks, 14 of them (40%) learnt to perform the feeding task, learning to associate the disk colour with the presence of food (Fig. 4). A different result was found in the control group were all the 8 observers were unable to learn by themselves how to solve the novel foraging task (Fig. 4). The difference between the two groups was significant (Fisher-Irwin exact test: p=0,039), confirming that observers used the observation of others to learnt the association task.

**Figure 4:**
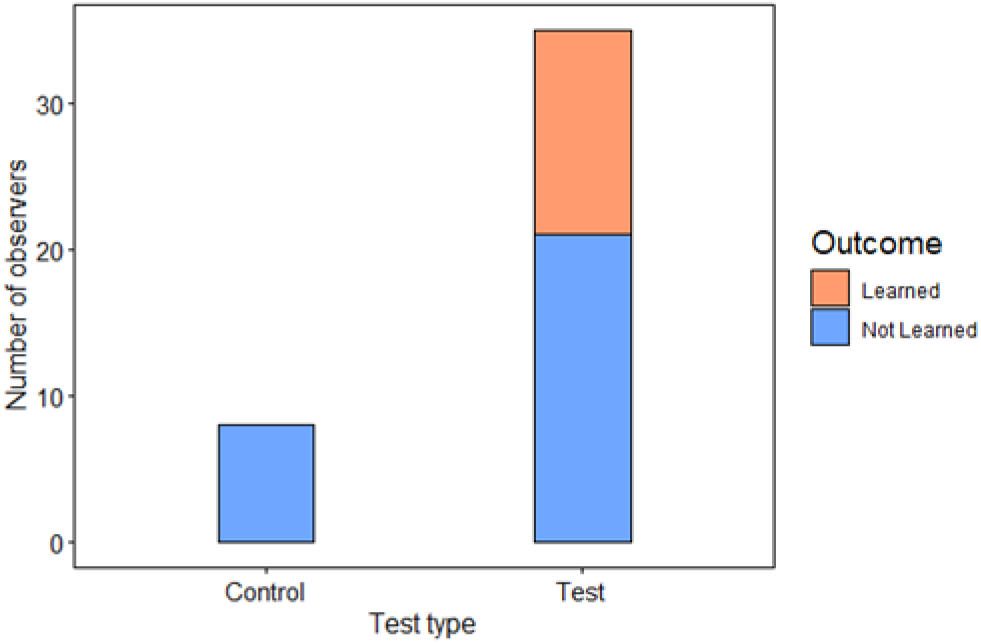
Outcome of the social learning test for the Control group (N=8) and the Test group (N=35).

The majority of observers (11 out of 14) took from 8 to 11 trials to learn (Average = 9). However, three of them (21%) were very quick to learn, having done it between the 3^rd^ and 5^th^ trials. So, there was some heterogeneity on how fast they were capable of learning. This could result from chance or from intrinsic differences between individuals.

We considered sex and age of both demonstrators (D_e_) and observers (O_b_) as possible explanatory factors for differences in learning ability. Learning was not significantly affected by either sex or age of demonstrators and observers or the interaction between them (Table 1). Bird morphology, in particular PC1, which accounts for wing length variation (Table A.1), was a significant predictor of social learning (B=0.969, p=0,038). That is, birds with longer wings were more capable of learning by observation. This difference in wing size is irrespective of birds age and sex, since these factors were included in the model. The GLM with PC1 as the explanatory variable had the lowest AICc and compared to all other models had a AICc delta larger than 2.

**Table 1:** Results of the Generalized Linear Model selection framework on the outcome of social learning test, comparing social learner and non-learners. Each row is the result of the GLM conducted with said predictor. Bold type indicates a significant result.

| Predictors | Estimate<br>d means | Standard<br>errors | B | *p* | Wald Chi-Square | AICc | Delta | Weight |
| --- | --- | --- | --- | --- | --- | --- | --- | --- |
| **PC1 Morph** | - | **0.4646** | **0.969** | **0.037** | **4.355** | **46.3** | - | **0.531** |
| Age Ob | 1Y 0.43<br>AD 0.71 | 0.7246 | -1.204 | 0.097 | 2.761 | 48.6 | 2.35 | 0.164 |
| Sex De | M 0.68<br>F 0.50 | 0.7026 | -0.773 | 0.271 | 1.211 | 50.3 | 3.98 | 0.072 |
| Sex Ob | M -0.69<br>F 0.53 | 0.7085 | -0.683 | 0.335 | 0.929 | 50.5 | 4.27 | 0.063 |
| Age De | 1Y 0.53<br>AD 0.69 | 0.7085 | -0.683 | 0.335 | 0.929 | 50.5 | 4.27 | 0.063 |
| PC2 Morph | - | 0.3207 | -0.147 | 0.648 | 0.209 | 51.1 | 4.79 | 0.048 |
| Age Ob*Age D | - | - | - | 0.498 | 2.375 | 51.1 | 4.81 | 0.048 |
| Sex Ob*Sex De | - | - | - | 0.202 | 4.622 | 53.9 | 7.65 | 0.012 |

### Memory and reversal learning

The Subgroup1 (n=9) was tested for memory retention 15 days after the ending of the social learning test. All of them succeeded in the foraging task (binomial test p=0.004), indicating that all birds retained in their memory the association they had learnt before.

Subsequently, the Subgroup1 was exposed to a reversal of the association rule, by placing the food under the other colour. Of the nine birds tested, 4 (44%) were capable of reversing the previously learnt rule. Age or sex did not explain the differences in reversal learning ability of the observers (both estimate = 0.2877; st. error=1.6073; p=0.858).

### Personality

#### Novel object test

Intra-individual repeatabilities were measured for each of the variables selected as proxy for neophobia/neophilia. *General Activity* and *Perch Near Object* were highly repeatable (R>0.5) (Table 2). The time *Perching on Feeder* was less repeatable and only marginally significant (R=0.302, P=0.049). We tested the variables for correlation (Table A.2), and we found that the variable *Perch Near Object* had a positive but marginally non-significant correlation with *Perch on Feeder* (0.477, P= 0.053), it also had a negative marginally non-significant correlation with the *General Activity* (−0.471, P= 0.056). This indicates that more active birds tend to be less prone to get close to the novel object. We found no significant correlation between *Perch on Feeder* and the *General Activity* (−0.205, P= 0.429). However, *General activity* was not significantly affected by the presence or absence of the novel object, so that it is not associated with boldness, contrary to the other two variables (Table A.2).

**Table 2:** Results of the repeatability test for each personality variable from the two personality tests. Type of distribution of the variables is indicated. R= Repeatability index; higher values indicate higher interindividual Repeatability. Bold type indicates high repeatability scores (R>0.5). In brackets are indicated the type of transformations applied to the respective variables.

| Variables | Test | Distribution | R | p |
| --- | --- | --- | --- | --- |
| Time Facing Mirror (sqrt) | Mirror | Normal | **0.589** | 0.0017 |
| Time Resting (Log_inv) | Mirror | Normal | **0.658** | 0.0004 |
| General Activity (Log) | Novel object | Normal | **0.504** | 0.0003 |
| Perch Near Object | Novel object | Binomial | **0.711** | 0.0010 |
| Perch on Feeder | Novel object | Binomial | 0.302 | 0.0486 |

#### Mirror test

In the mirror test there was high repeatability (R>0.5) for *Time Facing Mirror* and *Inverse of Time Resting* (Table 2), indicating individuals behaved consistently across the two test trials. The two variables were negatively correlated (−0.590, P=0.013), so that birds that spent more time facing the mirror were also those who spent more time resting (Table A.2). Both variables were significantly affected by the type of test (control or not), thus revealing to be capturing the sociability trait (Table A.2).

A correlation analysis between the variables of the Mirror Test and the Novel object test was not significant for all variables (Table A.2).

#### Personality and social learning

We performed two Principal Components Analyses (PCA) to produce indexes for novelty avoidance and for sociability. The Principal Component (PC) extracted from the Mirror test variables had positive loadings (0.892) on the *Inverse Time Resting* and negative (−0.892) on the *Time Facing Mirror* (Table A.3). Thus, in our Sociability Index more sociable individuals had a lower score. The PC obtained for the Novel Object test had a positive loading (0.946) on both latencies, *Perch Near Object* and *Perch on Feeder* (Table A.3). Thus, in our Boldness Index, a lower score meant a more curious individual. Both learners and non-learners have, on average, similar personality indexes. However, individuals that did not learn have a wider distribution of values for boldness than learners (Fig. 5a). Learners, have a wide distribution of sociability values (Fig. 5b). Neither the sociability index (p=0.994) nor the boldness index (p=0.364) were significant predictors of the differences in social learning ability.

**Figure 5:**
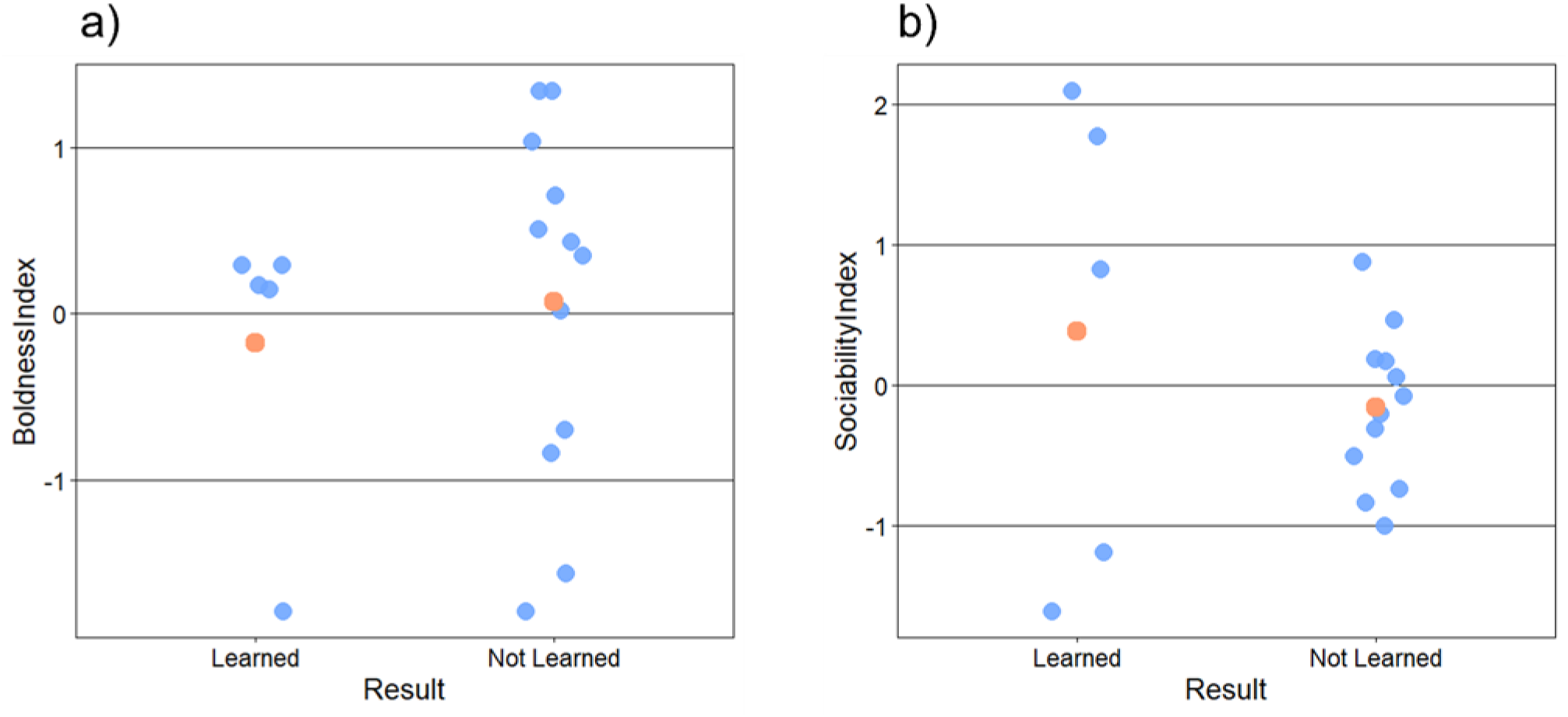
Scatter plots of the personality traits of successful and unsuccessful social learners. a) Boldness Index; b) Sociability Index. Orange dots represent mean values.

## DISCUSSION

We found evidence of social learning in the serin, a gregarious, granivorous small passerine bird. Individuals revealed considerable variation in their cognitive ability to deal with the challenge, with fewer than half having succeeded. Differences in learning ability did not appear to be dependent on age and sex of the individuals.

### Demonstrators training

We succeeded in training serins a colour-food association through a protocol with a gradual increase in the association difficulty (Aplin et al., 2013; Chantal et al., 2016) similarly to other studies that made use of foraging boards or similar (Ashton et al., 2018; Caves et al., 2018; Heyes and Saggerson, 2002). During the training the demonstrators took on average more attempts to learn in phase (A) than in the other phases (A=18.2; B=6.9; C=11.1), even though in principle this was the easiest step, because the food was visible. This learning pattern, and the fact that some individuals never interacted with the testing apparatus, suggests that these animals either do not pay attention or are intimidated by the experimental apparatus, refraining from exploring the available space, or do not understand the nature of the setup, that is, they can access food in some of the wells of the board.

### Social learning

In the last few decades many studies on social learning have broadened our understanding of this behaviour in an increasing number of animal species with different ecologies and life history traits (Hoppitt & Laland, 2013). This is the first study to investigate social learning of *Serinus serinus* and its interactions with personality and morphology. Here we showed that a granivorous and gregarious bird species, that is not an active hunter, such as the serin, significantly relies on social learning in a foraging context. Indeed, a relevant proportion (40%) of the individuals involved in the study was capable to learn a complex foraging technique involving associating colour-food in less than 300 minutes of exposure to a capable demonstrator. Although previous studies on sparrows have shown that a granivorous passerine can make use of social information in the foraging context (Katsnelson et al, 2011; Cadieu; Fryday et al, 1994), ours is the first study revealing the ability of granivorous birds to socially learn an associative rule.

We tested birds without a demonstrator in order to determine if they could have learnt the association colour-food just by trial and error, without using social information, as a control group. In order to make conditions more similar, they were also presented with other birds exploring the experiment, but which were naïve birds, that is, they could not provide information on the location of food. Neither the naïve birds were able to learn the associative task, nor the control observers, exposed to them. Thus, we could exclude trial and error or social facilitation as a possibility for the learning association between colour and food, confirming that the observers learnt by social learning.

One striking result from our experiment was that a large proportion of individuals was unable to learn the association rule, not making use of social information. This is not totally unusual, and in previous studies it was found that interindividual differences can affect reliance on social learning. In a few animal species, the sex of the observers is a reliable predictor of social learning. Female blue tits exposed to a demonstrator had higher learning success rate than males, with percentage of individual that learnt around 61% for females and 30% for males (Aplin et al., 2013). Similarly, females were faster social learners in two species of lemur (Kappeler 1987; Schnoell and Fichtel, 2012). In serins, we did not find any significant effect of sex on the learning ability of individuals, neither any interaction between the sexes of demonstrators and observers. Age is another factor that can affect social learning. Juveniles and young adults were the most successful social learners in lizards (Noble et al., 2014) and meerkats (Abrahms et al., 2021; Aplin et al., 2013; 2015 Thornton & Malapert 2009). While in birds contrasting results emerged from successive studies on blue tits, with the most recent proving no effect of age on learning (Penndorf & Aplin, 2019). In Serins, our results showed that young adults (57% of the successful individuals) did not learn more often than older birds.

Contrary to previous findings in great and blue tits (Teichmann et al., 2020) serins did not show a bias towards the red colour in the foraging context. Serins were equally likely to socially learn from a demonstrator trained to feed from red wells as well as one trained with blue wells.

We found a significant but weak association between learning and wing length. As far as we know it is the first time in the literature that a morphological trait is a significant predictor of social learning. The relationship involves some interpretation and, thus, this result has to be considered carefully and would benefit from further analysis. A possible explanation for this association is that better social learners are also more efficient foragers, so that they can invest more energy in developing longer wings. In fact, in this species, wings continue to grow during all adult life and not just during the early development (Mota, personal observation). This characteristic could support the idea that serins invest in wing length if they have enough resources to do so. Future studies could investigate these ideas.

### Cognitive skills

The cognitive skills of a subgroup of the successful observers were tested further. First, we wanted to know if there was memory retention of what was learnt. After 15 days all learnt serins still remembered the food-colour association, with all individuals succeeding in the foraging task. Proficient memory is a trait observed across many passerines from Corvidae (Tomback, 1980), and Paridae (Sherry, 1984), to Estrildidae (Ueno & Suzuki, 2014). However, our finding further extends this evidence to true finches (Fringillidae). This ability is useful to remember the distribution of food across the foraging area especially during the winter season.

We went further in testing more complex cognitive abilities of these birds, by subjecting those that learnt the association to a reversal leaning protocol. A fraction of those birds (44%) was able to reverse the associative rule previously learnt. Comparable results were obtained also in great tits, where 59 % of captive individuals succeeded in the reversal task, although not resulting from social learning (Cauchoix et al., 2017). Other studies on avian species found individuals to differ in reversal learning success or in reversal learning speed (Laschober et al., 2021; Zidar et al., 2018; Benedict et al., 2022). However, differences in the experimental designs makes comparisons scarcely meaningful, due to differences in the number of trials given to reach the learning criterion, and in the task to complete. The reversal learning ability revealed by serins has potential fitness consequences since cognitive plasticity is very important to cope with an everchanging environment. It is, thus, a skill that can be positively selected in these birds’ populations. While in some species females were found to perform better in reversal learning test (Petrazzini et al., 2017), we have no indication for a sex difference in serins, although the sample size is too small to draw conclusions.

### Personality

We tested serins for personality in order to determine whether personality differences could explain the differences in social learning that we were finding. There were strong repeatabilities on the personality traits tested. This constitutes a further support to the existence of stable behavioural differences related to personality in Serins, as a previously found (Leitão, 2011). We did not find any significant correlation between the variables of the Mirror Test and the Novel object test, hance boldness and sociability are probably not part of a personality syndrome. Although, our sample of 17 individuals limits our ability to have clear cut results, we found that boldness and sociability did not affect significantly the ability for social learning in serins. We predicted that bolder individuals would have an advantage in this learning setup because it involved the interaction with a novel object and environment, hence less neophobic individuals would have been less stressed during the tests. Besides, we expected more sociable individuals to be better social learners because they would interact and observe more the demonstrator they were paired with. Previous literature on the interactions between personality traits and social learning did not depict a clear pattern across the different species and testing conditions studied. The difference between the learning test protocols and the great diversity in ecology and life history traits of the studied species (from insects to birds, to primates) makes it extremely challenging to compare the results between studies on this topic. Even considering only the avian species studied, researchers found contrasting results: more explorative great tits performed better at social learning (Marchetti and Drent, 2000) while bolder and more explorative geese rely less on social information (Kurvers et al, 2010).

We selected a small number of personality tests, but that were adequate for the context of social learning. It is possible that other personality traits may be related with differences in learning ability, or maybe that ability has no relation to personality in any way.

## Conclusion

In conclusion our study showed that serins are capable of social learning in a foraging context. This study widens our knowledge on how widespread social learning is in non-social animals, investigating a new species in the field. It also shows that serins, and possibly other gregarious finches, can learn a stimulus-reward association through social learning, thus reducing the need for direct exposure. Serins not only learnt associative rules but also remembered them over a short period of time. Some individuals were even capable of reversal learning. Social learning abilities in Serins, as in other bird species, are extremely variable between individuals, even among those that belong to a single population. We investigated this interindividual difference and found no evidence of it being related to sex, age or personality. We found a weak positive relationship between social learning success and wing length; therefore, we hypothesize that social learning adaptiveness might increase foraging efficiency and thus translate in a higher amount of energy that the individual can invest in growing. To clarify this interaction a larger sample size and further studies on the actual impact of social learning on individual fitness would be required, including dominance. Serins behaved consistently throughout context and time with each individual exhibiting repeatable personality traits. However, these behavioral interindividual differences were not associated with individual ability for social learning.

## Appendix

**Table A.1:**
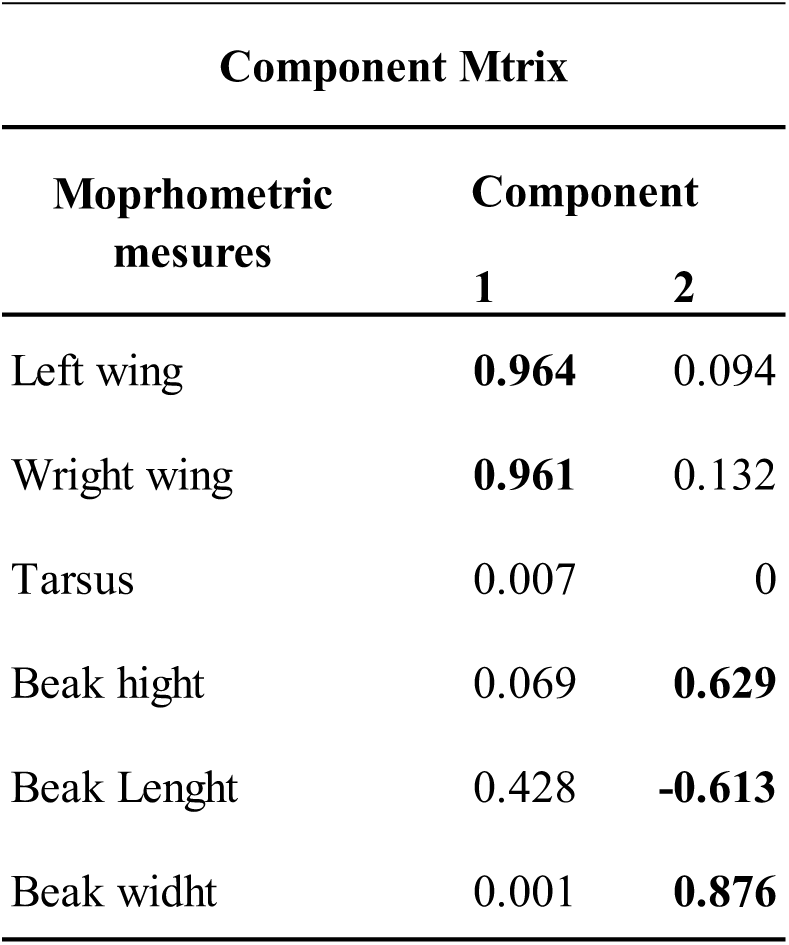
The component matrix of the Principal Component Analysis conducted on the morphometric variables. Two components are extracted, in bald type are the main loading for each component.

**Table A.2:**
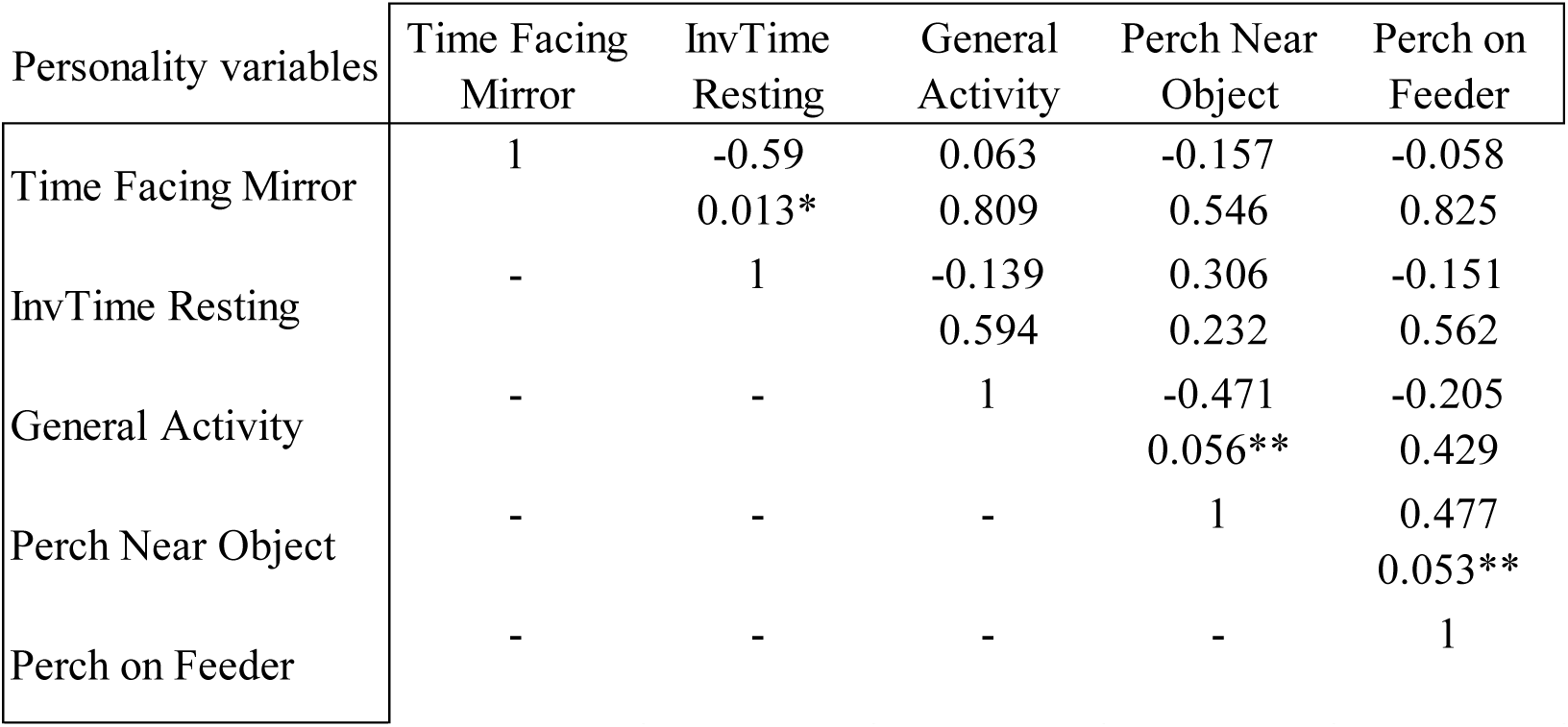
Pearson Correlation between the selected personality variables. In the first row is the Pearson correlation value, in the second row the significance (two-tailed). Asterisks indicate significant correlations with a 95% CI, double asterisks indicate significant correlations with a 90% CI.

| Personality variables | Time Facing<br>Mirror | InvTime<br>Resting | General<br>Activity | Perch Near<br>Object | Perch on<br>Feeder |
| --- | --- | --- | --- | --- | --- |
| Time Facing Mirror | 1 | -0.59<br>0.013* | 0.063<br>0.809 | -0.157<br>0.546 | -0.058<br>0.825 |
| InvTime Resting | - | 1 | -0.139<br>0.594 | 0.306<br>0.232 | -0.151<br>0.562 |
| General Activity | - | - | 1 | -0.471<br>0.056** | -0.205<br>0.429 |
| Perch Near Object | - | - | - | 1 | 0.477<br>0.053** |
| Perch on Feeder | - | - | - | - | 1 |

**Table A.3:** Component loading of the personality variables used to compute the personality indexes. Positive loadings indicate that the variable is positively correlated to the index, negative loadings indicate the opposite.

| Test | Variable | Component loading |
| --- | --- | --- |
| Novel object | Perch on Feeder | 0.946 |
| Novel object | Perch Near Object | 0.946 |
| Mirror test | Time Facing Mirror | -0.892 |
| Mirror test | Time Resting | 0.892 |

## References

1. Abrahms, B., Teitelbaum, C.S., Mueller, T. (2021). Ontogenetic shifts from social to experiential learning drive avian migration timing. Nat Commun 12, 7326. 10.1038/s41467-021-27626-5

2. Aplin, L. M., Farine, D. R., Morand-Ferron, J., Cockburn, A., Thornton, A., & Sheldon, B. C. (2015). Experimentally induced innovations lead to persistent culture via conformity in wild birds. Nature, 518(7540), 538–541. 10.1038/nature13998

3. Aplin, L. M., Sheldon, B. C., & Morand-Ferron, J. (2013). Milk bottles revisited: Social learning and individual variation in the blue tit, Cyanistes caeruleus. Animal Behaviour, 85(6), 1225–1232. 10.1016/j.anbehav.2013.03.009

4. Arbilly, M., & Laland, K. N. (2017). The magnitude of innovation and its evolution in social animals. Proceedings of the Royal Society B: Biological Sciences, 284(1848). 10.1098/rspb.2016.2385

5. Ashton, B. J., Ridley, A. R., Edwards, E. K., & Thornton, A. (2018). Cognitive performance is linked to group size and affects fitness in Australian magpies. Nature, 554(7692), 364–367. 10.1038/nature25503

6. Beecher, M. D., & Brenowitz, E. A. (2005). Functional aspects of song learning in songbirds. In Trends in Ecology and Evolution (Vol. 20, Issue 3, pp. 143–149). Elsevier Ltd. 10.1016/j.tree.2005.01.004

7. Benedict, L. M., Heinen, V. K., Sonnenberg, B. R., Bridge, E. S., & Pravosudov, V. v. (2023). Learning predictably changing spatial patterns across days in a food-caching bird. Animal Behaviour, 196, 55–81. 10.1016/j.anbehav.2022.11.005

8. Clare Mcw. H. Benskin, N.I. Mann, R.F. Lachlan, P.J.B. Slater. (2002). Social learning directs feeding preferences in the zebra finch, *Taeniopygia guttata*, Animal Behaviour, 64 (5), 823–828, 10.1006/anbe.2002.2005.

9. Bond, A. B., Kamil, A. C., & Balda, R. P. (2007). Serial Reversal Learning and the Evolution of Behavioral Flexibility in Three Species of North American Corvids (Gymnorhinus cyanocephalus, Nucifraga columbiana, Aphelocoma californica). Journal of Comparative Psychology, 121(4), 372–379. 10.1037/0735-7036.121.4.372

10. Brainard, M. S., & Doupe, A. J. (2002). What songbirds teach us about learning. Nature 2002 417:6886, 417(6886), 351–358. 10.1038/417351A

11. Bridges, A. D., Royka, A., Wilson, T., Lockwood, C., Richter, J., Juusola, M., & Chittka, L. (2024). Bumblebees socially learn behaviour too complex to innovate alone. Nature. 10.1038/s41586-024-07126-4

12. Cadieu, N., Fruchard, S., & Cadieu, J. C. (2010). Innovative individuals are not always the best demonstrators: Feeding innovation and social transmission in Serinus canaria. PLoS ONE, 5(1). 10.1371/journal.pone.0008841

13. Carter, A. J., Marshall, H. H., Heinsohn, R., & Cowlishaw, G. (2014). Personality predicts the propensity for social learning in a wild primate. PeerJ, 2014(1). 10.7717/peerj.283

14. Cauchoix, M., Hermer, E., Chaine, A. S., & Morand-Ferron, J. (2017). Cognition in the field: comparison of reversal learning performance in captive and wild passerines. Scientific Reports, 7(1), 12945. 10.1038/s41598-017-13179-5

15. Caves, E. M., Green, P. A., Zipple, M. N., Peters, S., Johnsen, S., & Nowicki, S. (2018). Categorical perception of colour signals in a songbird. Nature, 560(7718), 365–367. 10.1038/s41586-018-0377-7

16. Chantal, V., Gibelli, J., & Dubois, F. (2016). Male foraging efficiency, but not male problem-solving performance, influences female mating preferences in zebra finches. PeerJ, 2016(8). 10.7717/PEERJ.2409

17. Diana F. Tomback. (1980) How Nutcrackers Find Their Seed Stores, The Condor, Volume 82, Issue 1, Pages 10–19, https://doi-org.pros2.lib.unimi.it/10.2307/1366779

18. Díaz, M. (1994). Variability in seed size selection by granivorous passerines: effects of bird size, bird size variability, and ecological plasticity. Oecologia, 99, 1–6.

19. Dougherty, L. R., & Guillette, L. M. (2018). Linking personality and cognition: A meta-analysis. In Philosophical Transactions of the Royal Society B: Biological Sciences (Vol. 373, Issue 1756). Royal Society Publishing. 10.1098/rstb.2017.0282

20. Dubois, F., Giraldeau, L. A., & Réale, D. (2012). Frequency-dependent payoffs and sequential decision-making favour consistent tactic use. Proceedings of the Royal Society B: Biological Sciences, 279(1735), 1977–1985. 10.1098/rspb.2011.2342

21. Fiorito, G. & Scotto, P. (1992). Observational learning in Octopus vulgaris. Science n. 256.5056, pp. 545–547.

22. Fryday, & Greig-Smith, P. W. (1994). The Effects of Social Learning on the Food Choice of the House Sparrow (Passer domesticus). Behaviour, 128(3/4), 281–300. http://www.jstor.org/stable/4535177

23. Greene, E. (1987). Individuals in an osprey colony discriminate between high and low quality information. Nature, 329(6136), 239–241. 10.1038/329239a0

24. Grüter, C., Leadbeater, E., & Ratnieks, F. L. W. (2010). Social Learning: The Importance of Copying Others. Current Biology, 20(16), R683–R685. 10.1016/j.cub.2010.06.052

25. Grüter, C., & Ratnieks, F. L. W. (2011). Flower constancy in insect pollinators: Adaptive foraging behavior or cognitive limitation? In Communicative and Integrative Biology (Vol. 4, Issue 6, pp. 633–636). Landes Bioscience. 10.4161/cib.16972

26. Guillamón, A., Valencia, A., Calés, J., & Segovia, S. (1986). Effects of early postnatal gonadal steroids on the successive conditional discrimination reversal learning in the rat. Physiology & Behavior, 38(6), 845–849.

27. Ha, J. C., Mandell, D. J., & Gray, J. (2011). Two-item discrimination and Hamilton search learning in infant pigtailed macaque monkeys. Behavioural Processes, 86(1), 1–6. 10.1016/j.beproc.2010.07.010

28. Hämäläinen, L., Rowland, H. M., Mappes, J., & Thorogood, R. (2017). Can video playback provide social information for foraging blue tits? PeerJ, 2017(3). 10.7717/peerj.3062

29. Heyes, C. M. (1994). SOCIAL LEARNING IN ANIMALS : CATEGORIES AND MECHANISMS. In Biol. Rev (Vol. 69).

30. Heyes, C., & Saggerson, A. (2002). Testing for imitative and nonimitative social learning in the budgerigar using a two-object/two-action test. Animal Behaviour, 64(6), 851–859. 10.1006/ANBE.2003.2002

31. Hoppitt, W., & Laland, K. N. (2013). Social Learning: An Introduction to Mechanisms, Methods, and Models. Princeton University Press. http://www.jstor.org/stable/j.ctt2jc8mh

32. Hultsch, H., & Todt, D. (2004). Learning to sing. Nature’s Music: The Science of Birdsong, Elsevier, 80–107. 10.1016/B978-012473070-0/50006-2

33. Kameda, T., & Nakanishi, D. (2002). Cost–benefit analysis of social/cultural learning in a nonstationary uncertain environment: An evolutionary simulation and an experiment with human subjects. Evolution and Human. Behavior, 23(5), 373–393. 10.1016/S1090-5138(02)00101-0

34. Kappeler, P. M. (1987). The acquisition process of a novel behavior pattern in a group of ring-tailed lemurs (Lemur catta). Primates, 28(2), 225–228. 10.1007/BF02382571

35. Katsnelson Edith, Motro Uzi, Feldman Marcus W. and Lotem Arnon. (2011). Individual-learning ability predicts social-foraging strategy in house sparrows. Proc. R. Soc. B.278582–589 10.1098/rspb.2010.1151

36. Kendal, J., Giraldeau, L. A., & Laland, K. (2009). The evolution of social learning rules: Payoff-biased and frequency-dependent biased transmission. Journal of Theoretical Biology, 260(2), 210–219. 10.1016/j.jtbi.2009.05.029

37. Kimball, M. G., Gautreaux, E. B., Couvillion, K. E., Kelly, T. R., Stansberry, K. R., & Lattin, C. R. (2022). Novel objects alter immediate early gene expression globally for ZENK and regionally for c-Fos in neophobic and non-neophobic house sparrows. Behavioural Brain Research, 428. 10.1016/j.bbr.2022.113863

38. Kitowski, I. (2009). Social learning of hunting skills in juvenile marsh harriers Circus aeruginosus. Journal of Ethology, 27(3), 327–332. 10.1007/s10164-008-0123-y

39. Kurvers, R. H. J. M., van Oers, K., Nolet, B. A., Jonker, R. M., van Wieren, S. E., Prins, H. H. T., & Ydenberg, R. C. (2010). Personality predicts the use of social information. Ecology Letters, 13(7), 829–837. 10.1111/j.1461-0248.2010.01473.x

40. Laland, K. N. (2004). Social learning strategies. Animal Learning & Behavior, 32(1), 4–14. 10.3758/BF03196002

41. Laschober, M., Mundry, R., Huber, L., & Schwing, R. (2021). Kea (Nestor notabilis) show flexibility and individuality in within-session reversal learning tasks. Animal Cognition, 24(6), 1339–1351. 10.1007/s10071-021-01524-1

42. Leitão, A. M. V. (2011). Sexual selection: the influence of personality, behavioural and ornamental traits in the mate choice of Serin (Serinus serinus). Master thesis at University of Coimbra. https://eg.uc.pt/bitstream/10316/30822/1/TeseAnaLeitão2009101686.pdf

43. Marchetti, C., & Drent, P. J. (2000). Individual differences in the use of social information in foraging by captive great tits. Animal Behaviour, 60(1), 131–140. 10.1006/anbe.2000.1443

44. Mery, F., Varela, S. A. M., Danchin, É., Blanchet, S., Parejo, D., Coolen, I., & Wagner, R. H. (2009). Public Versus Personal Information for Mate Copying in an Invertebrate. Current Biology, 19(9), 730–734. 10.1016/j.cub.2009.02.064

45. Miletto Petrazzini, M. E., Bisazza, A., Agrillo, C., & Lucon-Xiccato, T. (2017). Sex differences in discrimination reversal learning in the guppy. Animal Cognition, 20(6), 1081–1091. 10.1007/s10071-017-1124-4

46. Noble, D. W. A., Byrne, R. W., & Whiting, M. J. (2014). Age-dependent social learning in a lizard. Biology Letters, 10(7). 10.1098/rsbl.2014.0430

47. Nomakuchi, S., Park, P. J., & Bell, M. A. (2009). Correlation between exploration activity and use of social information in three-spined sticklebacks. Behavioral Ecology, 20(2), 340–345. 10.1093/beheco/arp001

48. Pike, T. W., & Laland, K. N. (2010). Conformist learning in nine-spined sticklebacks’ foraging decisions. Biology Letters, 6(4), 466–468. 10.1098/rsbl.2009.1014

49. R Core Team (2023). R: A language and environment for statistical computing. R Foundation for Statistical Computing, Vienna, Austria. https://www.R-project.org/.

50. Réale, D., Dingemanse, N. J., Kazem, A. J. N., & Wright, J. (2010). Evolutionary and ecological approaches to the study of personality. In Philosophical Transactions of the Royal Society B: Biological Sciences (Vol. 365, Issue 1560, pp. 3937–3946). Royal Society. 10.1098/rstb.2010.0222

51. Rendell, L., Fogarty, L., Hoppitt, W. J. E., Morgan, T. J. H., Webster, M. M., & Laland, K. N. (2011). Cognitive culture: Theoretical and empirical insights into social learning strategies. In Trends in Cognitive Sciences (Vol. 15, Issue 2, pp. 68–76). 10.1016/j.tics.2010.12.002

52. Roelofs, S., Nordquist, R. E., & van der Staay, F. J. (2017). Female and male pigs’ performance in a spatial holeboard and judgment bias task. Applied Animal Behaviour Science, 191, 5–16. 10.1016/j.applanim.2017.01.016

53. Rosa, P., Nguyen, V., & Dubois, F. (2012). Individual differences in sampling behaviour predict social information use in zebra finches. Behavioral Ecology and Sociobiology, 66(9), 1259–1265. 10.1007/s00265-012-1379-3

54. Schnoell, A. V., & Fichtel, C. (2012). Wild redfronted lemurs (Eulemur rufifrons) use social information to learn new foraging techniques. Animal Cognition, 15(4), 505–516. 10.1007/s10071-012-0477-y

55. Sherry, D. (1984). Food storage by black-capped chickadees: Memory for the location and contents of caches. Animal Behaviour, 32(2), 451–464. 10.1016/S0003-3472(84)80281-X

56. Sih, A., & Del Giudice, M. (2012). Linking behavioural syndromes and cognition: A behavioural ecology perspective. In Philosophical Transactions of the Royal Society B: Biological Sciences (Vol. 367, Issue 1603, pp. 2762–2772). Royal Society. 10.1098/rstb.2012.0216

57. Slagsvold, T., & Wiebe, K. L. (2011). Social learning in birds and its role in shaping a foraging niche. In Philosophical Transactions of the Royal Society B: Biological Sciences (Vol. 366, Issue 1567, pp. 969–977). Royal Society. 10.1098/rstb.2010.0343

58. Smit, J. A. H., & van Oers, K. (2019). Personality types vary in their personal and social information use. Animal Behaviour, 151, 185–193. 10.1016/j.anbehav.2019.02.002

59. Stoffel, M. A., Nakagawa, S., & Schielzeth, H. (2017). rptR: repeatability estimation and variance decomposition by generalized linear mixed-effects models. In Methods in Ecology and Evolution (Vol. 8, Issue 11, pp. 1639–1644). British Ecological Society. 10.1111/2041-210X.12797

60. Stoffel, M. A., Nakagawa, S., & Schielzeth, H. (2019). An introduction to repeatability estimation with rptR. CRAN. R-Project. Org. Retrieved from https://cran.rproject.org/web/packages/rptR/vignettes/rptR.html

61. ’Svensson, L. (1992). Identification Guide to European Passerines (British Trust for Ornithology, Ed.; 4th ed.). British Trust for Ornithology.

62. Teichmann, M., Thorogood, R., & Hämäläinen, L. (2020). Seeing red? Colour biases of foraging birds are context dependent. Animal Cognition, 23(5), 1007–1018.

63. Templeton, J. J., Luc, ·, & Giraldeau, A. (1996). Vicarious sampling: the use of personal and public information by starlings foraging in a simple patchy environment. In Behav Ecol Sociobiol (Vol. 38).

64. Thornton, A., & Malapert, A. (2009). Experimental evidence for social transmission of food acquisition techniques in wild meerkats. Animal Behaviour, 78(2), 255–264. 10.1016/j.anbehav.2009.04.021

65. Thornton, A., & Samson, J. (2012). Innovative problem solving in wild meerkats. Animal Behaviour, 83(6), 1459–1468. 10.1016/j.anbehav.2012.03.018

66. Trompf, L., & Brown, C. (2014). Personality affects learning and trade-offs between private and social information in guppies, Poecilia reticulata. Animal Behaviour, 88, 99–106. 10.1016/j.anbehav.2013.11.022

67. Ueno, A. and Suzuki, K. (2014). Visual Learning and Retention in Birds. Animal Science Journal, 85: 186–192. https://doi-org.pros2.lib.unimi.it/10.1111/asj.12092

68. Valera, F., Wagner, R. H., Romero-Pujante, M., Gutiérrez, J. E., & Rey, P. J. (2005). Dietary specialization on high protein seeds by adult and nestling serins. The Condor, 107(1), 29–40.

69. Van Bergen, Y., Coolen, I., & Laland, K. N. (2004). Nine-spined sticklebacks exploit the most reliable source when public and private information conflict. Proceedings of the Royal Society B: Biological Sciences, 271(1542), 957–962. 10.1098/rspb.2004.2684

70. Zidar, J., Balogh, A., Favati, A., Jensen, P., Leimar, O., Sorato, E., & Løvlie, H. (2018). The relationship between learning speed and personality is age- and task-dependent in red junglefowl. Behavioral Ecology and Sociobiology, 72(10). 10.1007/s00265-018-2579-2

